# Single-strain behavior predicts responses to environmental pH and osmolality in the gut microbiota

**DOI:** 10.1101/2022.08.31.505752

**Authors:** Katharine M. Ng, Sagar Pannu, Sijie Liu, Juan C. Burckhardt, Thad Hughes, William Van Treuren, Jen Nguyen, Kisa Naqvi, Bachviet Nguyen, Charlotte A. Clayton, Deanna M. Pepin, Samuel R. Collins, Carolina Tropini

## Abstract

Changes to gut environmental factors such as pH and osmolality due to disease or drugs correlate with major shifts in microbiome composition; however, we currently cannot predict which species can tolerate such changes or how the community will be affected. Here, we assessed the growth of 92 representative human gut bacterial strains spanning 28 families across multiple pH values and osmolalities *in vitro*. The ability to grow in extreme pH or osmolality conditions correlated with the availability of known stress response genes in many cases, but not all, indicating that novel pathways may participate in protecting against acid or osmotic stresses. Machine learning analysis uncovered genes or subsystems that are predictive of differential tolerance in either acid or osmotic stress. For osmotic stress, we corroborated the increased abundance of these genes *in vivo* during osmotic perturbation. The growth of specific taxa in limiting conditions in isolation *in vitro* correlated with survival in complex communities *in vitro* and in an *in vivo* mouse model of diet-induced intestinal acidification. Our data show that *in vitro* stress tolerance results are generalizable and that physical parameters may supersede interspecies interactions in determining the relative abundance of community members. Importantly, we provide an extensive resource for predicting shifts in microbial composition and gene abundance in complex perturbations. Furthermore, this work highlights the physical environment as a major driver of bacterial composition and the importance of performing physical measurements in animal and clinical studies to elucidate the drivers of shifts in microbiota abundance.

**Significance Statement:** Changes in pH and particle concentration (osmolality) commonly result from gut disease or the ingestion of common drugs, causing changes in bacterial growth and microbiota composition within the intestine. Thus far, the effects of physical parameters on the growth of intestinal bacterial taxa have not been well documented in the context of predicting microbiota community composition. To address this gap, we examined the growth of 92 bacterial species under varying pH and osmolality conditions. We found that physical parameters are key predictors of bacterial abundance in individual-strain cultures and in complex bacterial communities. Moreover, our results identified specific genes and pathways that are predictive of growth in specific environments. Together, these findings can aid in determining the effectiveness of microbiota therapies in gut environments subjected to various perturbations.

## Introduction

The animal digestive tract naturally consists of numerous distinct environments in which physical and chemical conditions such as oxygen concentration, acidity, mucosal stiffness, and temperature are tightly regulated by host–microbial interactions. Environmental gradients along the intestine create a continuum of habitats that the microbiota explores in its voyage along the intestinal tract (1–3). Alterations in pH and particle concentration (osmolality) commonly occur in gut disease or result from the ingestion of specific compounds. For example, inflammatory bowel disease (IBD), intestinal cancers, and antacids are associated with abnormal pH values (4, 5). Furthermore, diarrhea and aging are associated with increased oxygen in the intestine, and conditions such as malabsorption due to intolerances (e.g., celiac disease) or overconsumption of salts, alcohol, and laxatives lead to changes in osmolality (6–8). Additionally, the microbiota affects physical parameters by consuming luminal oxygen, degrading the mucosal layer, or acidifying the environment through fermentation and short-chain fatty acid (SCFA) production (9–11).

Changes in the gut physical environment affect the gut microbiota on a broad scale, favoring growth only when the biochemical and physical conditions match the requirements of each taxon (3). The survival of specific taxa is driven by the function of genes and pathways that regulate metabolism and stress responses over short and chronic time scales. These genes and pathways play a key role in establishing the microbiota members that can grow in specific regions of the gut and host states (1–3). For example, the steep intraluminal oxygen gradient partitions strictly anaerobic bacteria such as *Faecalibacterium* away from the more oxygenated epithelium, while more aerotolerant bacteria such as Enterobacteriaceae can associate with the mucosa (12). Beyond oxygen sensitivity, pH and osmolality also impact bacterial growth and survival (4, 13–15). Even small alterations in pH and osmolality can dramatically affect bacterial growth due to alterations in enzyme activity, energetic favorability of certain nutrient substrates, and rates of protein synthesis (16–18).

Previous studies have highlighted broad differences among microbial taxa in their adaptation to pH and osmolality, as evidenced by the differential enrichment of well-studied taxa in response to pH alterations (13). For example, *Lactobacillus* species propagate over wide ranges of pH, whereas acidic environments inhibit some members of the *Bacteroides* genus (19). Despite these general trends, certain members within these taxonomic groups excel in high-stress conditions of high pH and osmolality, while others display sensitivity to these parameters, indicating genus- or even strain-specific differences (4, 20). Even in the absence of limiting cases in which taxa cannot grow, changing physical conditions can affect bacterial growth rates, resulting in microbiota composition shifts within the highly competitive and nutrient-depleted environment of the intestine. For example, mild osmotic diarrhea induced by polyethylene glycol (PEG) can induce long-term changes in gut microbial membership despite presenting no change in bacterial density or load (14).

Thus far, the effects of physical parameters on bacterial growth across intestinal bacterial taxa have not been well documented. Importantly, the taxonomic level at which growth phenotypes can be generalized remains unclear; moreover, it remains unknown whether the physical environment is broadly predictive of bacterial response and abundance. Closing this gap of understanding is particularly critical for identifying the relationship between the microbiota and disease, as the presence of certain gut bacterial members may be strongly dictated by the physical environment rather than disease-specific phenotypes. In addition, identifying the genes that allow particular gut members to survive varying physical parameters is crucial. This knowledge would also shed light on the effectiveness of microbiota therapies, as procedures such as fecal microbiota transplant or probiotic administration may be rendered ineffective by the transfer of members that cannot survive in the disease-altered environment.

In this work, we examined the growth phenotypes of 92 species from 28 families across a range of pH and osmolality values. We combined high-throughput growth measurements, environmental measurements, and machine learning-assisted comparative genomics to systematically identify the capacity of microbial taxa to survive in pH and osmolality conditions relevant to health and disease. We performed a thorough *in vitro* analysis of bacterial growth of individually grown strains and revealed general trends of tolerance among phylogenetically related microbes, including known pathogens and probiotics strains. We corroborated these results by performing human microbiota community experiments *in vitro* and in humanized mouse models and demonstrated the broad predictability of bacterial abundance across multiple donors and conditions. Importantly, we found that *in vitro* results of single-strain stress tolerance can predict bacterial behavior in complex *in vivo* conditions. We also found that the presence of genes involved in osmotic stress response is predictive of survival in disrupted osmolality environment. Together, our results demonstrate that the physical environment is broadly predictive of bacterial response and abundance. This knowledge will aid in determining the effectiveness of microbiota therapies and in assessing whether therapies will be viable in a given perturbed gut environment.

## Results

### Collection of 92 strains from 28 families of bacteria

We cultured 92 bacterial strains from 28 common gut bacterial families across seven phyla, comprising a diverse set of strains with a focus on human isolates (**Figure 1A**). We chose these strains based on their public availability, fully sequenced genomes, and broad interest due to their prevalence in the gut microbiota. We derived most strains from the BEI collection of Human Microbiome Project human strains, the American Type Culture Collection (ATCC), the Deutsche Sammlung von Mikroorganismen und Zellkulturen (DSMZ), and the Collection of Inflammation-Associated Mouse Intestinal Bacteria (CIAMIB) (21) (Materials and Methods). Of these strains, 69 were human isolates, 10 were mouse-derived, and the remaining 13 were either isolated from probiotics or other strain types. Some characterized probiotic strains (11) were isolated from commercial sources and were not previously sequenced. For taxa of high interest due to their prevalence, abundance, or health relevance, we included multiple species or strains within a family in order to avoid drawing species-specific conclusions (22–30). Specifically, we increased the coverage of the Bacteroidaceae, Bifidobacteriaceae, Lactobacillaceae, Lachnospiraceae, Enterobacteriaceae, and Prevotellaceae families. To facilitate high-throughput cultivation and comparisons of strains, we grew the majority of the bacteria (83/92) in anaerobic conditions in Mega Medium, a rich and undefined medium previously demonstrated to support the growth of a wide variety of strains (Materials and Methods) (31). The remaining strains required more specialized media for growth (Materials and Methods, **Figure 1B**, Table S1). To measure the impact of bacterial growth on the environmental pH, we supplemented the experimental media with 2’,7’-bis-(2-carboxyethyl)-5-(and-6)-carboxyfluorescein (BCECF), which enabled real-time pH measurements coupled with optical density (OD) measurements (**Figure 1B**). We uncovered no significant trends in medium acidification with respect to growth in different conditions (**Figure S1A**). As the genomes in our strain library have been fully sequenced and assembled, we performed comparative genomics analyses that combined protein annotations from the Pathosystems Resource Integration Center (PATRIC). To visualize this analysis across all strains, we created a novel visualization tool to explore PATRIC annotation data and compare across multiple genomes (https://tropinilab.shinyapps.io/strain_heatmap_app/) (**Figure 1C**). We grew the strains in eight different conditions, spanning four osmolalities (from unmodified medium osmolalities of 0.23–0.44 Osm/kg to a maximum condition of 1.8 Osm/kg) and four pH values (4–7.4). We selected these ranges to mimic the potential environmental conditions that intestinal bacteria encounter along the gastrointestinal tract and during perturbations (14, 32, 33).

**Figure 1.**
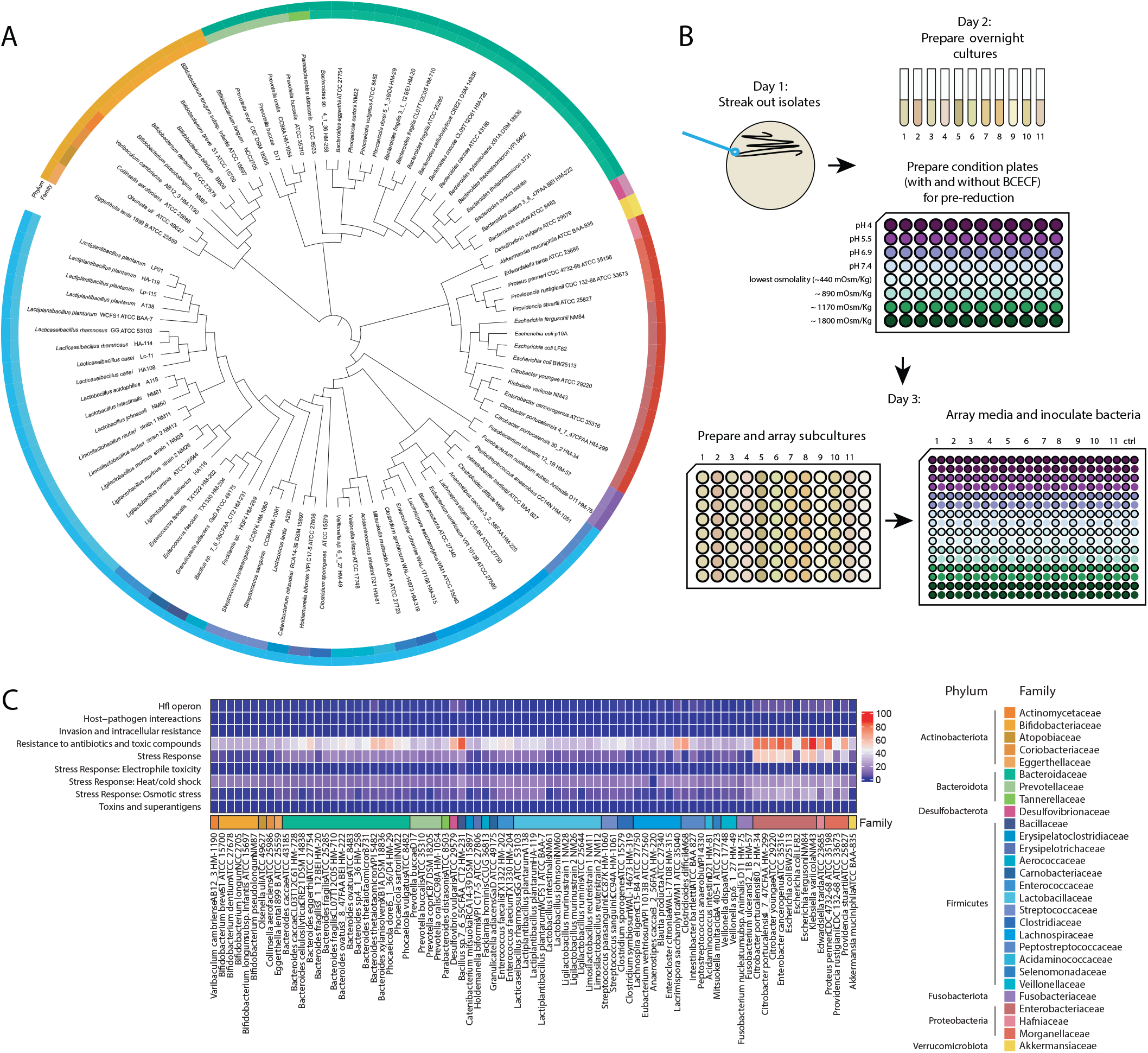
Phylogenetic overview and experimental setup of characterized intestinal bacterial strains. **(A)** 16S rRNA sequences from each member of the strain library were acquired from the SILVA database and used to generate a phylogenetic tree. **(B)** Experimental design and workflow for the characterization of growth of bacterial strains under different physical conditions. **(C)** Heatmap of PATRIC annotations of characterized strains within the subcategories of the Stress Response, Defense, and Virulence gene categories.

### Bacterial families display a range of tolerance to increasing osmolality

Increasing osmolality elicited widely divergent effects on the bacterial families assayed in this study (**Figure 2A**). While *in vivo* measurements of high gut osmolality are usually less than ~1100 mOsm/kg (14), we sought to explore a wider range of high osmolalities, as many bacterial taxa showed strong growth at these values in the current study. Lactobacillaceae and Enterobacteriaceae family members displayed robust growth at moderately high (~1176 mOsm/kg) and high (~1800 mOsm/kg) osmolalities. Interestingly, we observed modest heterogeneity among genera and species within Lactobacillaceae; the strains that were more negatively affected in high-osmolality conditions were *Lactobacillus murinus* strains 1 (NM26) and 2 (NM28), *Lactobacillus intestinalis* NM61, and *Lacticaseibacillus rhamnosus* HA-114. Conversely, bacteria in the Bacteroidaceae, Bifidobacteriaceae, and Lachnospiraceae families displayed a wide range of sensitivities to increasing osmolality; except for two *Bifidobacterium* strains, these bacteria were unable to grow in media with an osmolality of ~1800 mOsm/kg. Multiple species were extremely sensitive to osmolality, including the mucin degrader *Akkermansia muciniphila* ATCC BAA-835, most Prevotellaceae species tested, and Erysipelotrichaceae member *Holdemanella biformis* VPI C17-5 ATCC 27806. While many bacterial strains showed decreased growth rates under high osmolality, they still reached maximum yields similar to those achieved in normal osmolality, indicating the same ability to leverage nutrients in these limiting conditions (**Figure S1B**).

**Figure 2.**
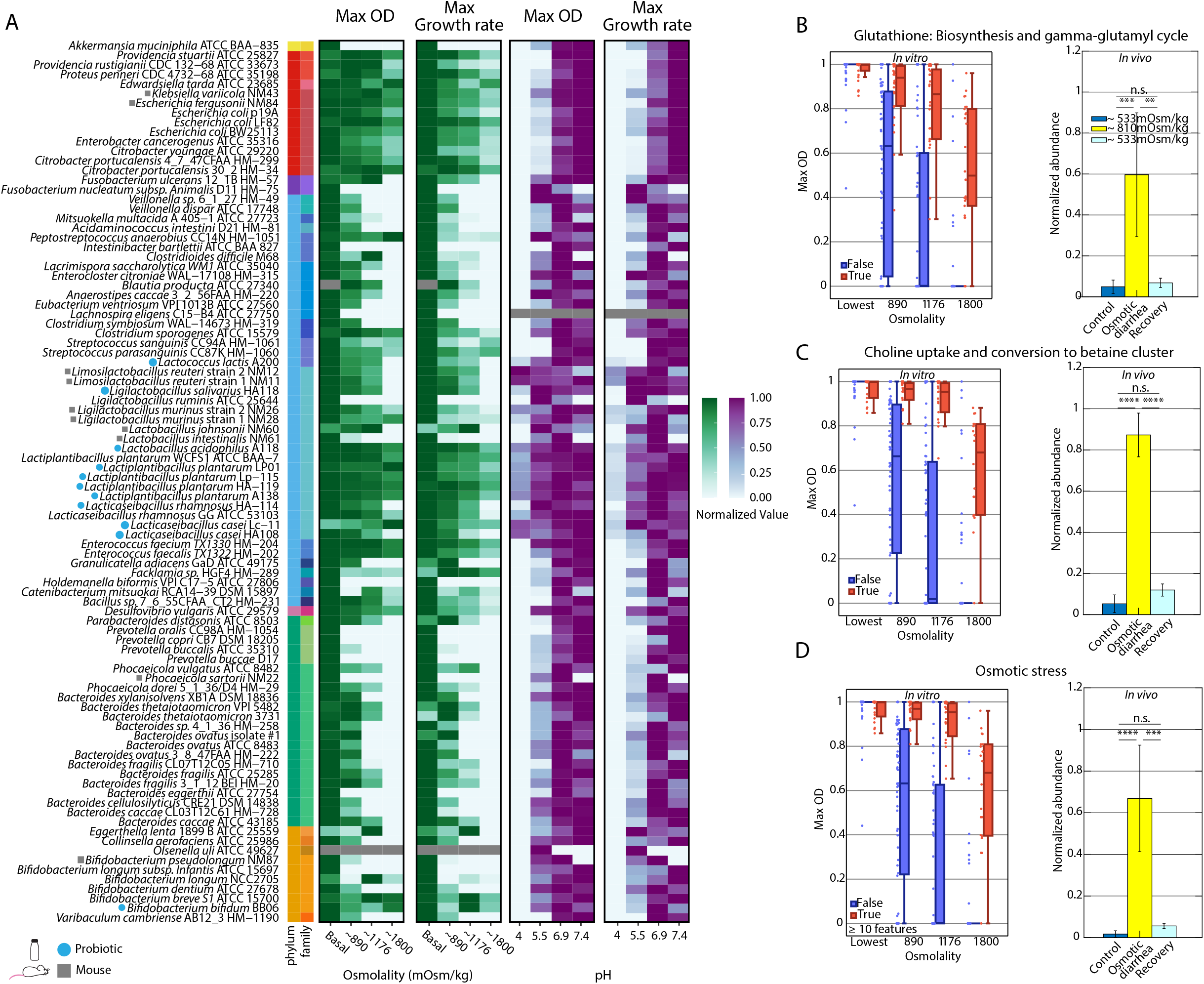
Osmotic and pH stress responses lead to phenotype variations. **(A)** Heatmap displaying normalized growth rate and OD_600_ of 92 characterized bacterial strains in osmotic conditions ordered by phylogenetic relatedness. The growth rates and OD were normalized to maximum growth rate and OD values within a characterized strain across osmolality conditions. The growth curves characterized in conditions of varying osmolality and pH demonstrate general trends of tolerance across bacterial taxa. **(B–D)** ML analysis of PATRIC annotations and growth profile data of characterized strains obtained via a one-level decision tree regression model. **(B)** Boxplot of the presence (red) or absence (blue) of bacterial species possessing features in glutathione biosynthesis and the gamma-glutamyl cycle as a function of the maximum OD across osmolality conditions of ~440, 890, 1176, 1800 mOsm/kg (left). This pathway is found to be enriched *in vivo* during osmotic perturbation (right). **(C)** Boxplot of characterized bacterial species encoding (red) or missing (blue) a gene for the choline uptake and conversion to betaine cluster as a function of the maximum OD under varying osmolality conditions (left). This pathway is found to be enriched *in vivo* during osmotic perturbation (right). **(D)** Boxplot highlighting the presence (red) or absence (blue) of at least 10 osmotic stress genes as a function of the maximum OD in varying osmotic conditions (left). This pathway is found to be enriched *in vivo* during osmotic perturbation (right). ** p <0.01, *** p <0.001, **** p < 0.0001

Having created a resource of growth abilities for a broad set of gut bacteria, we sought to discover the mechanisms that underlie the tolerance of some bacteria to different osmotic conditions by finding genes or functions that are consistently over-represented in tolerant bacteria. As the majority of genomes (81/92) in our strain library have been fully sequenced and assembled, we performed comparative genomics analyses by combining genomic annotations conducted using tools created by PATRIC with our quantified growth phenotypes (34).

Next, we employed a machine learning (ML) strategy to determine whether potential annotations employed by PATRIC were indicative of a strain’s ability to grow in varying environmental osmolality and pH conditions. Unlike most ML applications, where the goal is to train a model based on input features and then evaluate and optimize its accuracy and generalization, our goal was restricted to the task of feature selection from the large set of available PATRIC annotations. Therefore, we constructed a novel featurization of the PATRIC annotations and used a simple ML model called a decision stump to fit many predictive models to the data. For each candidate PATRIC feature, we measured modeling error on both a train set and held-out test set. We then used the modeling error to rank all the PATRIC features according to their ability to predict the phenotype. This ML analysis efficiently identified PATRIC annotations (ML model features) that correlated with an increased maximum OD of bacterial growth in the different growth conditions (Materials and Methods, Tables S2, S3).

We found that the presence of several subsystems correlated with a higher maximum OD at high osmolality (Table S2). A challenge with this type of analysis is that many functional annotations may be correlated and present in the same strains (**Figure S2**), but not necessarily directly implicated in response to perturbations. Thus, we identified features that ranked high in distinguishing growth phenotypes and were mechanistically plausible. Importantly, the held-out dataset performed comparably to the training dataset, indicating the identified features generalized across this sampling of strains (**Figure S3A**). To further increase our ability to detect relevant features, we re-analyzed a previously published and annotated metagenomic dataset of *in vivo* osmotic perturbation in a humanized mouse model (14). Briefly, in this dataset, mice were exposed to the osmotic laxative PEG, which increased the mean intestinal osmolality from 533 to 810 mOsm/kg. The functional pathways present in this community were then quantified for mice prior to, during, and after osmotic perturbation. We identified features that were both over-represented in high osmolality *in vivo* and detected in our ML analysis.

Of particular interest were the subclasses for glutathione biosynthesis/gamma-glutamyl cycle, choline uptake and conversion to betaine, and osmotic stress (**Figures 2B–D**). Notably, we also identified cold shock proteins as a distinguishing feature for growth in different osmotic conditions (**Figure S3B,C**). Characterized bacterial taxa that possessed annotated genes within these subsystems or roles demonstrated a higher maximum OD on average with increasing osmolality and were also significantly over-represented *in vivo* in high-osmolality conditions. Confirming the importance of stress gene annotations, we identified the osmotic stress gene category as the top predictive feature in this analysis, with a minimum of 10 features in this category being predictive of growth in high-osmolality conditions (**Figure 2D**, Table S3).

### Bacterial families demonstrate a wide range of phenotypic variation in growth and yield in response to acidic/alkaline stress

We assessed the growth of the strain library for a pH range of 4–7.4 and revealed a wide range of tolerances to acidic conditions (**Figure 2A**). Because physiological pH within the gut varies with intestinal site, diet, and disease, we chose a range of pH conditions to encompass physiologically relevant perturbations a bacterial species may face (4, 22, 23, 25). Most strains, except for Lactobacillaceae members, were unable to grow at pH 4; even at pH 5.5, these strains displayed serious defects in growth rate and maximum yield. Bacteria within the Lactobacillaceae family displayed the highest tolerance to low pH, as expected for lactic acid bacteria. However, even within this family, genera differed in their tolerance, with several members of *Lactobacillus* and *Ligilactobacillus* displaying sharp decreases in growth rate and maximum OD at pH 4 and inhibition of growth at pH 7.4 compared with physiological pH (6.4-7). Interestingly, isolates from commercial probiotic sources showed high sensitivity to pH changes. Furthermore, at pH 5.5, many species, such as Erysipelotrichaceae members, displayed a decreased growth rate and yield. An exception were members of the Desulfovibrionaceae, Fusobacteriaceae, Veillonellaceae, and Bifidobacteriaceae families, which contain members that have been described in acidic environments (e.g., acidic mine tailings, dental caries, vagina, and fermented foods, respectively) (35–39). Reports have also shown that *Bacteroides* species, which belong to the family Bacteroidaceae, are sensitive to low pH (40); however, we observed a wide range of sensitivities in growth at pH 5.5 in this genus, suggesting that different species and strains may be better adapted to acidic conditions. Some families displayed severe deficits in growth rate and yield at pH 5.5, including members of the Enterobacteriaceae and Streptococcaceae families. An acid-tolerance response has been described for members of the Enterobacteriaceae family (e.g., *Escherichia coli* and *Salmonella*) (41); however, this response may require priming in mildly acidic conditions prior to exposure to more acidic conditions. In our experiments, we subcultured bacteria at neutral pH immediately prior to growth under experimental conditions to simulate the transfer of a healthy microbiota into a diseased environment, which could potentially mask acid-stress adaptations in these families. Interestingly, several bacteria displayed narrow pH tolerances and were inhibited by mildly alkaline conditions, including Lactobacillaceae members. This sensitivity to alkaline conditions has been documented for Lactobacillaceae (19). Interestingly, similar to our observations in high-osmolality conditions, relative deficits in growth rate did not always translate into deficits in final yield (**Figure 2A**), suggesting a path for survival of species in communities experiencing physical perturbation in the gut, if they are able to withstand washout.

After identifying growth patterns across our strain library, we once again performed ML analysis on PATRIC features to identify subsystems correlated to bacterial strain growth in acidic/alkaline stress. Unlike our osmolality analysis, the identified PATRIC features were sensitive, in that the model’s fit of the held-out data showed poorer generalization (Table S3, **Figure S4**). Furthermore, PATRIC features ranked as highly predictive strongly correlated with specific bacterial families such as Lactobacillaceae, and therefore had less broadly predictive power across the identified strains.

### Taxon-specific responses to pH and osmolality are predictive of behavior in naturally derived complex communities

In the intestine, the microbial communities comprising the microbiota are affected by the physical environment as well as other microbial species that compete for resources and may produce inhibitory molecules (42). To determine whether the behaviors in single-strain pure cultures are generalizable to growth phenotypes in communities, we examined the growth of complex gut microbiota in *in vitro* cultures subjected to defined pH and osmolality environments. We cultured feces from six healthy human donors for 48 hours in Mega Medium under the same medium conditions used to assess the single-strain growth of the individual strains. Selection of fecal microbiota from multiple unrelated donors enabled us to study different taxa that are naturally coexisting and adapted to their specific complex community, allowing us to identify generalizable behaviors independent of specific metabolic interactions. Varying pH and osmolality resulted in a wide range of community compositions (**Figure 3A**). We compared abundance changes in different conditions at the family level, avoiding potential strain-specific behaviors. We observed numerous and distinct positive and negative correlations between pH and osmolality and the relative abundance of specific families (**Figure 3B**). A negative correlation with pH indicates that a family is more tolerant of low pH, while a negative correlation with osmolality indicates that a family is less tolerant of high osmolality.

**Figure 3.**
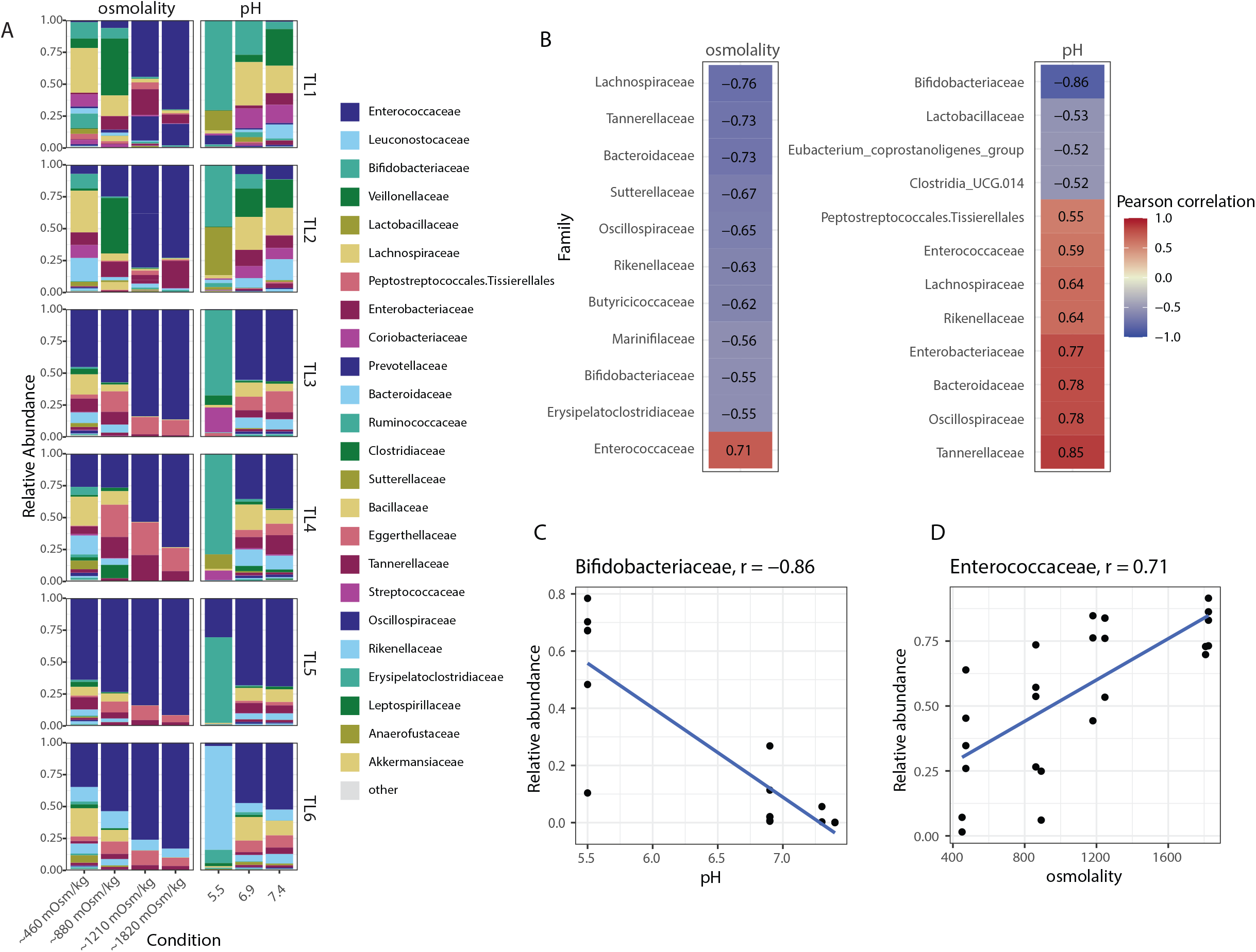
The relative abundance of bacterial families correlates with osmolality and pH across multiple complex microbiota communities *in vitro*. **(A)** Bar plot of relative abundance of bacterial taxa isolated from *in vitro* cultures of human fecal samples (n = 6) subjected to ranges of pH and osmolalities identical to *in vitro* single-strain cultures. The relative abundance and composition of bacterial families were determined via 16S rRNA sequencing. **(B)** Pearson’s correlations of bacterial families in complex communities of human fecal samples characterized *in vitro*, with negative correlations between relative abundance and osmolality (left) and pH (right) highlighted in blue and positive correlations in red. **(C)** Plot of relative abundance as a function of increasing pH for Bifidobacteriaceae, demonstrating a negative correlation between pH and relative abundance (r = −0.86). **(D)** Plot of relative abundance and increasing osmolality for Enterococcaceae, demonstrating a positive correlation between osmolality and Enterococcaceae abundance (r = 0.71).

These correlations of human isolates for varying pH mirrored our observations in the single-strain growth of individual species, suggesting that, for specific taxa, the environmental pH is a stronger driver of bacterial abundance than the specific microbiota composition. For example, the relative abundance of Bifidobacteriaceae exhibited a highly negative correlation (r = −0.86) with increasing pH (**Figure 3C**), matching the widespread low pH tolerance of Bifidobacteriaceae species relative to other families (**Figure 2A**). In contrast, the Tannerellaceae, Oscillospiraceae, and Bacteroidaceae families correlated positively with increasing pH (suggesting acid sensitivity), which is consistent with our single-strain growth data.

Responses of bacterial families to different osmolalities in fecal fermentations also displayed similarities to single-strain responses. For example, the relative abundance of Enterococcaceae exhibited a highly positive correlation with osmolality (r = 0.71) (**Figure 3D**), in line with the tolerance of Enterococcaceae species to high osmolality *in vitro* (**Figure 2A**) and *in vivo* (14). Conversely, members of the Bacteroidaceae family displayed a strong negative correlation with osmolality; the heterogeneity of single-strain responses in this family **(Figure 2A)** suggests that the Bacteroidaceae population within the surveyed communities may be skewed toward relatively osmolality-sensitive members.

For many strains, the single-strain behavior was predictive of their response in a community setting; however, we observed a weak correlation with environmental pH or osmolality for some bacterial families whose members in our strain library displayed consistent responses *in vitro*. For example, some correlations for bacterial species that demonstrated acid tolerance in *in vitro* single-strain growth (e.g., Lactobacillaceae) were not as strong as expected, suggesting that other factors such as nutritional or resource competition may contribute to the relative abundance of bacterial species in the intestine (**Figure 3B**). Additionally, the Enterobacteriaceae family displayed increased abundance in increasing osmolality in only some fecal communities (Tropini Lab 1 [TL1], TL2, and TL4), which may be due to genus- or species/strain-level variation or, more likely, out-competition by other osmotically tolerant bacteria present in non-responsive fecal communities. Because we used 16S ribosomal RNA (rRNA) amplicon sequencing and measured relative abundances, the survival and proliferation of other osmotically tolerant bacteria could mask survival or increases in absolute Enterobacteriaceae abundance. We also observed a heterogeneity in responses across donors, which likely stems from differences in the presence or absence of bacterial families in individual donor samples. For example, Bifidobacteriaceae were the dominant family at low pH in all samples except for one (TL6), in which the lactic acid bacterial family Leuconostocaceae dominated; this family was absent or present at less than 1% in all samples except for TL6.

### Diet influences bacterial taxa composition in a manner predicted by in vitro pH and osmolality resilience

Having shown the *in vitro* generalizability of pH and osmolality resilience in a complex microbiota for several key taxa, we explored whether these phenotypes would also be consistent *in vivo*, where, beyond microbiota interactions, the interplay with host dynamics plays a significant role in determining bacterial abundance. As other studies have investigated the response to osmolality *in vivo* (14, 43), we sought to investigate how the microbiota is affected by changes in pH *in vivo*. We reasoned that changing diet would impact gut pH differentially in the various intestinal compartments (43). Specifically, microbial fermentation of carbohydrates in the cecum and colon produces SCFAs, lowering the pH of these intestinal compartments. As with any perturbation *in vivo*, changing diet will have multiple orthogonal effects (i.e., in this case, combining differences in nutrient availability for both the host and the microbiota as well as pH); however, using our *in vitro* analysis, we reasoned this model might enable us to identify patterns that are consistent with pH tolerance. Although most monosaccharides and disaccharides are hydrolyzed and absorbed in the small intestine, many dietary oligosaccharides and polysaccharides cannot be hydrolyzed by host enzymes and pass undigested into the large intestine, where they are readily fermented by bacteria. One such carbohydrate is guar gum (44), a galactomannan polysaccharide comprised of a linear backbone of β 1,4-linked mannose residues with randomly attached β-1,6-linked galactose residues, a structure that cannot be digested by the mammalian host (45). We hypothesized that mice on this diet would undergo increased fermentation and acidification in their large intestines relative to mice on a standard diet. We gavaged germ-free Swiss Webster mice with feces from a healthy human donor, colonizing their intestines with a representative community of human intestinal bacteria (**Figure 4A**). After equilibration for 6 weeks on a standard rodent diet, the mice were divided into two groups: one group was placed on a diet with 30% guar gum and allowed to adjust to their diet for an additional 2 weeks whereas the second group continued on a standard rodent diet (**Figure 4A**). We then sacrificed the mice and performed 16S rRNA sequencing (**Figure 4B**) and pH measurements (**Figure 4C**) on different intestinal segments. Measurements of intestinal content revealed a significant decrease in pH in the jejunum, cecum, and colon of mice receiving the guar gum diet (**Figure 4C**), suggesting that increased fermentation occurred. Unlike the pH at the other sites, the duodenal and ileal pH values were unaffected by the diet change. We then investigated whether SCFA production in the cecum was altered by the diet. Indeed, we found that butyrate levels increased three-fold in mice fed the guar gum diet while other SCFAs were not significantly affected (**Figure 4D**).

**Figure 4.**
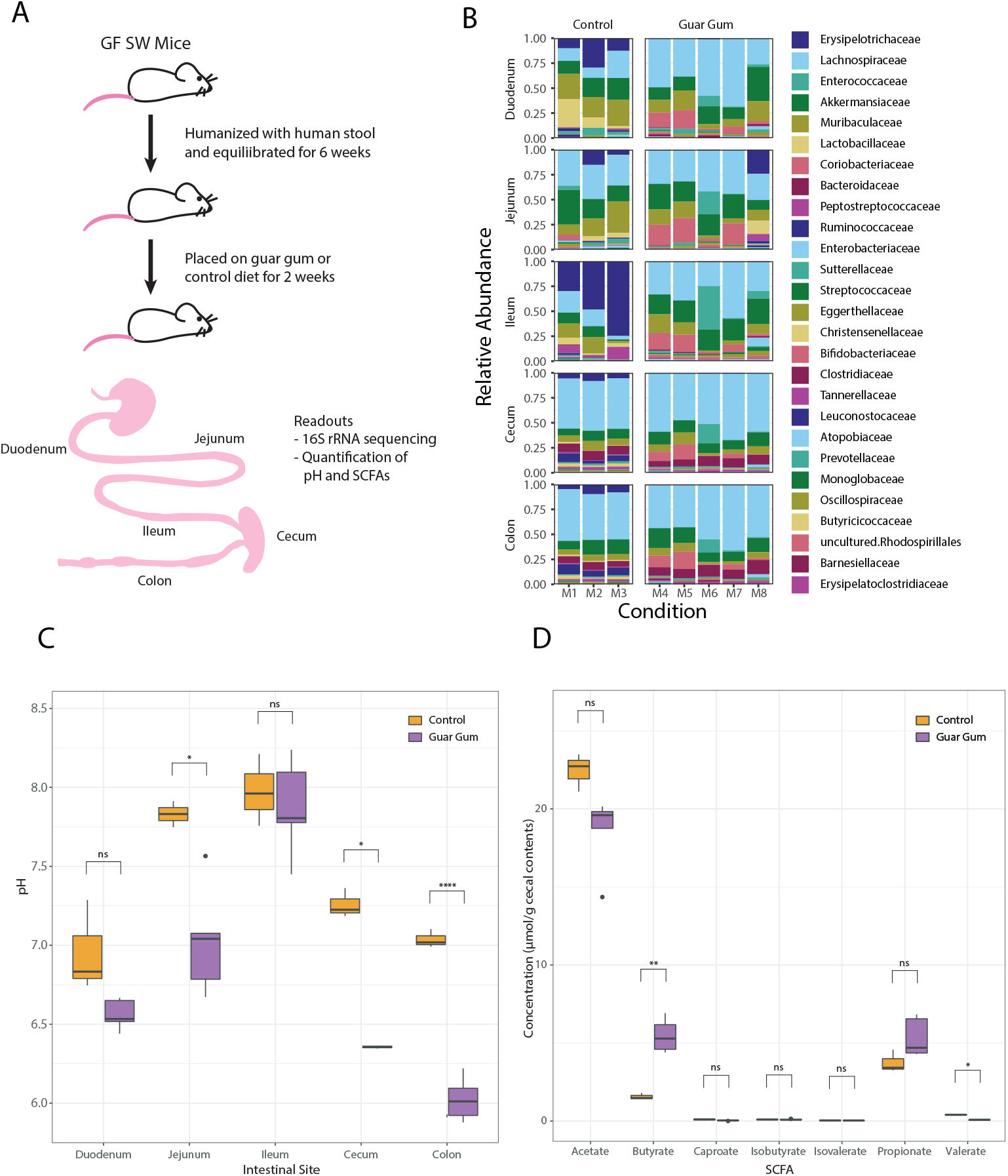
Humanized mice on a guar gum diet demonstrate a significant drop in cecal pH and shifts in bacterial family composition. **(A)** Experimental schematic of germ-free Swiss Webster (SW) mice, detailing the timeline of humanization and the switch to a guar gum diet (n = 5) versus a control diet (n = 3). The bottom figure depicts segments of the gastrointestinal tract collected for pH measurements, 16S sequencing, and SCFA measurements. **(B)** Relative bacterial abundance of humanized mice on the guar gum or control diet, highlighting the gut microbial composition at the family taxonomic level. **(C)** Quantification of pH along the gastrointestinal tract of humanized mice on the guar gum or control diet. **(D)** SCFA concentrations in cecal contents from humanized mice on the guar gum or control diet.

Next, we analyzed the relative abundance of bacteria in the different intestinal segments and diet conditions by performing 16S rRNA sequencing of communities isolated from those regions. In mice on the guar gum diet, the average cecal and colonic pH values were 6.36 and 6.02, respectively, compared with 7.26 and 7.04 for the standard-diet-fed mice. Given the distinct pH tolerance profiles of different bacterial families (**Figure 2A**), we hypothesized that this acidification would change the composition of the colonic microbiota. We observed an increase in the butyrate producer *Blautia* and a loss of the Erysipelotrichaceae family (**Figure 4B**). Our single-strain analyses revealed that *Blautia* growth was not inhibited at pH 5.5, consistent with its ability to thrive in these conditions *in vivo* (**Figures 2A & 4B**). Conversely, our *in vitro* data suggest that many *Bacteroides* members are sensitive to low pH (**Figure 2A**). Interestingly, we observed *in vivo* expansion of *Bacteroides* within the cecum and colon in experimental mice despite *Bacteroides* members displaying a sensitivity to *in vitro* conditions of pH 5.5 and below (**Figures 2A & 4B**). This observation suggests that the expansion of relatively more acid-tolerant *Bacteroides* isolates within the same niche may occur upon changes in the physical environment.

## Discussion

In this study, we sought to characterize the pH and osmolality tolerance of a wide range of publicly available and sequenced gut bacteria and probiotic strains. We characterized the growth of 92 bacterial representatives of the gut across 28 families under multiple pH and osmolality conditions *in vitro* (**Figure 1A**). Most of these bacterial species were human isolates, selected to maximize their relevance to the human intestine; however, by measuring the response to pH and osmolality in both mouse and human isolates from the same families, we provided a more extensive coverage of the diversity found within gut bacteria across multiple hosts. We developed high-throughput growth assays to test bacterial growth in pH and osmolality conditions representative of those found within the gut in health and disease. Our measurements demonstrate a wide range of tolerance to perturbation across bacterial families and family-specific responses to changes in pH and osmolality. Within families, individual members across hosts demonstrated a varied response to *in vitro* conditions. Using comparative genomics, we uncovered the abundance and prevalence of genes in characterized microbial taxa responding to stress. In many cases, the abilities to tolerate acidic stress and high osmolality were congruent with the abundance and presence of identified stress response genes. One limitation of this study arises from differences in the depth of gene annotations/knowledge of the more deeply studied families, such as Enterobacteriaceae and Bacteroidaceae, and more newly discovered and relatively culture-intractable families (**Figure 2A**). However, our analysis provides a framework for identifying strains that may possess novel stress responses for cases in which an isolate shows growth in limiting pH or osmolality conditions despite not possessing annotated stress tolerance genes.

We observed widespread pH and osmolality tolerance in representatives of Lactobacillaceae (including isolates from commercially available probiotic sources) and Enterococcaceae, respectively. Conversely, Bacteroidaceae and Bifidobacteriaceae displayed heterogeneity in their responses to osmolality and pH. Despite a broad diversity of growth phenotypes, we did not observe genomic features that could explain the phenotypic variation. This may be due to a difference in unannotated genes, or in the expression of annotated genes, and suggests that transcriptomic analyses or single gene knockout libraries may be required to untangle mechanisms underlying the differential tolerance of these bacteria.

Interestingly, many strains that displayed deficits in maximum growth rate at lower pH or higher osmolality still produced similar maximum yields (**Figure S1B**). Although bacteria in the intestine must grow at a sufficient rate to prevent washout, maintaining a maximum efficiency of biomass yield may also be an effective strategy for survival; thus, efforts to connect single-strain tolerance to behavior/yield in complex communities must incorporate both metrics. It is also important to note that while some of these strains displayed relative deficits in growth rate compared with their maximal growth rate in ideal conditions, a strain merely needs to survive or efficiently produce biomass relative to other bacteria to be propagated in the gut. Moreover, some bacterial taxa spatially inhabit specific niches within the gut, which limits the competition for resources against other bacteria preventing washout.

Our ML analysis highlighted how the presence of specific genes is strongly predictive of growth at different osmolalities (**Figures 2B–D**). Many of these genes are involved in stress tolerance (46, 47). For osmotic stress, the subsystem of glutathione biosynthesis/gamma-glutamyl cycle has previously been shown to provide osmo-adaptation (46). Tellingly, mutants lacking genes within this subsystem (e.g., *gshA* and *gor*) in *E.coli* show deficient growth in elevated osmolarity (46). Furthermore, we identified other genes not traditionally viewed as osmotic-stress response genes (Table S2). For example, we found that dihydrolipoamide dehydrogenase is predictive of osmotic tolerance. This protein functions to oxidize dihydrolipoamide in a ping-pong mechanism as an oxidoreductase (48). Interestingly, the identified dihydrolipoamide dehydrogenase of the pyruvate dehydrogenase complex has been implicated in the increased osmo-tolerance of *Staphylococcus aureus* (49, 50).

The identification of genes and pathways involved in pH tolerance was less obvious than that for osmolality, indicating that family-specific tolerance mechanisms may be at play, masking potential generalizable features. This finding indicates that community-based techniques such as metagenomics may not shed light onto the importance of specific genes involved in pH tolerance; identified features that may be unrelated to pH may be over-represented in acid-tolerant bacteria and appear significant. Therefore, we predict that more traditional genetic screens and transcriptomics assays will be needed to discover genes involved in pH tolerance in poorly annotated bacteria. For cases in which there is phenotypic variation within a family, comparative genomic techniques may be valuable, but for cases in which there is complete penetration of a phenotype within a taxon, elucidation of such features will prove more challenging. Importantly, we also found that the most acid- and osmolality-tolerant bacteria generally did not overlap (**Figure 2**), suggesting that there are distinct mechanisms for acid and osmolality tolerance. This was also confirmed in our ML analysis, in which the set of features that predicted osmolality tolerance did not rank highly in pH tolerance (**Figure S5**).

We explored whether single-taxon phenotypes are generalizable to a complex microbiota grown in complex media (**Figure 3**). Using six distinct human fecal microbiota samples, we tested growth across pH and osmolality conditions and found that growth patterns observed in isolated bacteria were consistent in complex microbiota communities. Bifidobacteriaceae and Bacteroidaceae were negatively and positively correlated with pH, respectively, in concordance with their acid tolerance and sensitivity in single-strain growth (**Figure 2A**). Similarly, families such as Enterococcaceae exhibited positive correlations with osmolality, mirroring their tolerance to osmolality. Enterobacteriaceae also displayed osmotic tolerance in pure cultures and in three donor samples (**Figure 3**, TL1, TL2, and TL4); this family flourished in intermediate and high osmolalities as well. The lack of significant correlation across donors may be due to several reasons, including genetic or phenotypic variation at the species level and out-competition by other osmo-tolerant bacteria such as Enterococcaceae. Similarly, we also observed a bloom in the lactic acid bacteria family Leuconostocaceae in one donor sample (**Figure 3**, TL6) at pH 5.5; this species, which is found in fermented food products, was absent in three donors and present at approximately 0.1% or less in TL3 and TL5, where it appeared that Bifidobacteriaceae dominated in low pH conditions. This result underscores the importance of quantifying tolerance across multiple bacterial families to capture the potential diverse responses in heterogenous human populations.

Finally, we investigated whether our *in vitro* observations were representative of an *in vivo* microbiota responding to changes in gut pH (**Figure 4**). We selected an animal model with a dietary intervention that yielded a decrease in pH in multiple intestinal segments and a significant increase in butyrate, a SCFA found in cecal contents. These changes corresponded to a loss of Erysipelotrichaceae, a family that displayed extreme sensitivity to pH in single-strain growth with member *Holdemanella biformis* (**Figure 2A**). Additionally, our *in vitro* strain characterizations confirmed the *in vivo* phenotypes we previously observed in an animal model of mild osmotic diarrhea induced by PEG laxatives (14). In our *in vitro* strain library, the families Enterobacteriaceae and Enterococcaceae displayed high osmotic tolerance; these families experienced significant expansions during PEG treatment in humanized mice (14). Similarly, the family Verrucomicrobiaceae (which contains the species *Akkermansia muciniphila*) was extremely sensitive to osmolality in our *in vitro* characterization; this species decreased 25-fold during osmotic perturbation *in vivo* (14).

Together, these results highlight the importance of the physical parameters of pH and osmolality and their role in the survivability of bacterial taxa found within the gut and, therefore, overall gut community composition. By quantifying the pH and osmolality tolerance across a wide range of representative intestinal bacterial families, we found that *in vitro* tolerance to physical parameters in single-strain growth can predict the effect of changes on complex communities in an *in vivo* physical environment. The tolerances we investigated were also consistent in *in vivo* animal models for multiple taxa. Thus, quantifying taxon-specific responses of the gut microbiota to environmental perturbations provides key information regarding the dynamics of community changes during health and disease.

Beyond the consistency between individual and community growth patterns in different environmental conditions, a valuable implication of our studies is the importance of broadly characterizing the physical environment in microbiota studies. To better understand how to remediate diseases, such as IBD, associated with both dysbiosis and environmental perturbation of the gut, it is crucial to establish the physical parameter ranges in healthy and perturbed environments, including the microenvironments along the intestine that are relevant to disease states. Once these environmental parameters have been quantified, efforts can be made to predict how the existing pH and osmolality may affect the survival of prospective communities in the gut. These measurements are also important in microbiota therapies, where the tolerance of probiotic strains to osmotic and acidic perturbations must be identified to determine survivability and potential function as a therapeutic within a perturbed gut environment. Potential probiotic strains must be able to propagate and compete against resident microbes in an environment to provide therapeutic effects. Our work has demonstrated that probiotics within a family may differ in their tolerance to physical parameters; we observed strong heterogeneity among Lactobacillaceae (**Figure 2A**), which contains many currently marketed probiotic strains. For a diseased environment in which physical parameters are altered or misregulated, our results suggest these probiotics may not fare equally well. Thus, the selection of a particular strain must consider physical measurements from the disease state of interest as well as the tolerance of potential therapeutic probiotics.

Overall, these results indicate that the physical environment is a key predictor of bacterial abundance over a broad range of conditions and across multiple communities. This predictability across physiological ranges highlights the importance of monitoring the physical environment in microbiota studies as a key driver of bacterial availability and the utility of determining the diverse individual responses of bacteria in single-strain cultures.

## Materials and Methods

### Phylogenetic tree construction

We acquired 16S sequences for most bacterial species from the SILVA database (https://www.arb-silva.de/search/) and the National Center for Biotechnology Information (NCBI) (Project ID: PRJNA474907). Sequences downloaded from SILVA were at least 1500 bp in sequence length. The downloaded FASTA files were compiled into a single file and imported to MEGA 11.0.10: Molecular Evolutionary Genetics Analysis version 11 for alignment using the MUSCLE algorithm and for construction of a phylogenetic tree using the “Construct/Test Neighbour-Joining Tree” option (53, 54). We then uploaded the Newick file generated by MEGA 11.0.10 to iTOL v6 (https://itol.embl.de/) for modification and coloring (51).

### Bacterial culture

The bacterial strains and corresponding metadata (i.e., taxonomy) used in this study are reported in Table S1. All bacterial strains were grown and inoculated in a vinyl anaerobic chamber (Coy Laboratories) maintained with an atmosphere of 5% CO_2_, 5% H_2_, and 90% N_2_ (Linde Canada). All strains were incubated at 37°C for growth, and all glycerol stocks were stored at −80°C.

### Bacterial media

We prepared Mega Medium using the protocol provided in the Supplementary Methods, with minimal modifications from a previous publication (31). Each batch of liquid and solid media was autoclaved and pre-reduced in an anaerobic chamber at least 24 hours before use. To characterize the pH and osmolality tolerance of strains, we aseptically loaded liquid Mega Medium into two sterile 96-deep-well plates, with media adjusted to eight different conditions. Medium conditions consisted of Mega Medium adjusted to pH 4, 5.5, 6.9, or 8 (osmolality normalized to ~600 mOsm/kg, the osmolality of the pH 4 condition) or to osmolality conditions of ~440, ~890, ~1176, or ~1800 mOsm/kg. We adjusted medium osmolality conditions using sodium chloride. Lowest osmolality condition of adjusted media was dependent on the basal osmolality of media used to characterize strains ranging from (~234 mOsm/kg to ~440 mOsm/kg). We adjusted the medium pH using 10N HCl and NaOH (33% wt. solution in water). For pH measurements, we calibrated a micro pH probe (Orion™ PerpHecT™ ROSS™ Combination pH Micro Electrode, Cat. No. 8220BNWP) for media adjustment. For osmolality measurements, we injected 20 μL of media into an Advanced Instruments Osmo1® Single-Sample Micro-Osmometer using an Ease-Eject 20-Microliter Sampler and clean sampler tips. We filter-sterilized the media using 150-mL 4-μm vacuum filtration tops (VWR: 10040-444) after adjustment. Due to the relatively high base osmolality of the pH media, some strains could not grow in any pH conditions and were therefore grayed out in the heatmap (**Figure 2A**). We loaded one plate with media containing 1 μg/mL BCECF (Thermo Fisher) to measure the environmental pH during growth. BCECF is a fluorescent pH sensor that detects extracellular changes in medium pH by establishing a pH-dependent ratio of emission intensity at excitation wavelengths of 440 and 490 nm. We loaded another plate without BCECF to accurately determine the BCECF signal during pH calculations. Both plates were incubated in an anaerobic chamber for 24 hours to pre-reduce and equilibrate the media for anaerobic growth. For some strains that were unable to grow in Mega Medium, we used solid and liquid media of peptone yeast glucose (PYG) medium, brain heart infusion-supplemented (BHIS) medium + mucin, and Mega Medium supplemented with either lactate or a combination of sodium citrate and MgSO4, as noted in Table S1.

### Bacterial growth

Bacterial isolates were streaked onto solid media to isolate individual colonies from glycerol stocks and were grown at 37°C; single colonies were picked after 24–48 hours and cultured in “overnight” pre-reduced liquid media at 37°C for 16–36 hours. We primarily used Mega Medium for strain characterization, with a few exceptions (Table S1). Overnight cultures were diluted 10-fold in pre-reduced liquid media and incubated at 37°C for 2 hours. We then diluted the cultures 5-fold into pre-reduced liquid media in a 96-well plate in preparation for high-throughput strain loading. Subsequently, we added 5 μL of cultures to 75 μL of each medium condition in a 384-well plate, resulting in a 16-fold dilution with an 80-fold dilution in total. Each 384-well plate consisted of eight conditions varying in pH and osmolality, with each strain grown in quadruplicate in each condition. Three of these replicates contained BCECF, and one replicate was BCECF-free, which enabled real-time environmental pH measurements coupled with OD measurements. Cultures grew at 37°C, and we measured the absorbance at 600 nm and BCECF fluorescence every 13 minutes using a Synergy H1 plate reader (BioTek) for 48–96 hours of growth. We measured BCECF fluorescence using excitation wavelengths of 440 and 490 nm detected at 535 nm for every time point on the growth curve. To quantify the medium pH, we determined the ratio of emission intensity between excitation at 490 nm versus 440 nm and calibrated this ratio to a calibration curve of pH values measured before each experiment, according to Invitrogen’s protocol. We conducted our analysis using a custom-made MATLAB program (https://github.com/Tropini-lab/Strain_library_paper).

### Growth analysis

Growth curves were run through a custom MATLAB script (https://github.com/Tropini-lab/Strain_library_paper). Briefly, the program identifies replicates based on assigned metadata (strain, pH/osmolality, etc.) and automatically selects the three most similar replicates for each condition for averaging and plotting. The maximum growth rate for each OD curve is determined after a least-squares fit is performed for the OD curve to the Gompertz equation (52).

### Growth data standards

We selected appropriate growth data for strains grown in each condition by comparing growth data against control conditions. Controls consisted of sterile Mega Medium (or BHIS + mucin, PYG, or Mega Medium supplemented with lactate or MgSO4 + sodium citrate); strains that increased in OD at the same time as or after control wells were discarded and re-run in subsequent experiments. For bacterial strains that were selected and considered clean, we performed outlier detection on the quadruplicate OD measurements and selected the best three out of four technical replicates in each condition for downstream analysis.

### Isolation of commercial probiotics for characterization

We dissolved probiotics purchased from local pharmacies in sterile 1X phosphate-buffered saline (PBS; Fisher Bioreagents: BP3991) in an anaerobic chamber. The dissolved slurry was streaked onto agar plates and incubated anaerobically at 37°C for 24 hours. Both PBS and medium were pre-reduced in an anaerobic chamber for at least 24 hours before use. We isolated Lactobacillaceae using Mega Medium and Bifidobacteriaceae using *Bifidobacterium* selective iodoacetate mupirocin medium according to a previously published method (Table S1) (53).

### Stock preparation

We obtained bacterial isolates from multiple culture collections, including BEI, ATCC, and DSMZ. Source cultures were streaked onto Mega Medium agar or appropriate media as noted in Table S1, and single colonies were picked and frozen for storage using a 1:1 mixture of culture and a 50% glycerol solution. The solid and liquid rich media used for stock production are listed in Table S1. We confirmed the purity of final cultures via Sanger sequencing of the 16S rRNA gene using 8F and 1391R primers (8F: 5’-AGAGTTTGATCCTGGCTCAG-3’, 1391R: 5’-GACGGGCGGTGWGTRCA-3’).

### PATRIC annotations

Genomes of publicly available species were downloaded from NCBI and submitted to PATRIC (https://www.patricbrc.org/) for annotation (54). The NCBI taxonomy ID and domain (i.e., bacteria) are required for submission. The abundance of annotated genomic features was compared across species at subsystem levels using the Shiny library in RStudio.

### In vitro growth and PATRIC subsystem analysis

Analyses and graphing were performed using R V4.1.2 and RStudio V1.4.1717. We conducted heatmap analyses of RAST (rapid annotation using subsystem technology) subsystems and growth data through an in-house-developed R library named “strains_heatmaps” (available for download in the following GitHub repository: (https://github.com/Tropini-lab/Strain_library_paper) and the ComplexHeatmaps package v2.10.0 (55, 56). In brief, our R library assembles all tables downloaded from RAST into a data frame that compares the number of features present in the different strains. Then, the R library filters and collapses the data frame based on broad annotation categories (in our case, subsystems involved in acid/osmolality tolerance) and transforms the data frame into a format compatible for use with the ComplexHeatmaps library. The ComplexHeatmaps library is then implemented to make various heatmaps, with coloring based on feature counts for each subsystem for each strain and the growth data joined as additional heatmaps or heatmap annotations. For more detailed explanations, please refer to the scripts and tutorials in the GitHub repositories (https://github.com/Tropini-lab/Strain_library_paper).

### Machine learning (ML)

The goal of our ML model was to determine which PATRIC annotations could predict a strain’s ability to grow in varying pH and osmolality conditions.

#### Model input feature preparation

Using Python (version 3.10.5), we constructed a tabular Pandas (version 1.4.3) DataFrame of features for each strain’s genome starting from the PATRIC subsystem annotation output (57). PATRIC maps the strain genome name (*genome_name* in Table S4) to many PATRIC IDs, each of which is annotated with a *Superclass, Class, Subclass, Subsystem* name and *Role ID*. Because this is a one-to-many mapping of *genome_name* to PATRIC ID, each *genome_name* appears on multiple rows, with various numbers of PATRIC IDs for each *genome_name*.

To efficiently fit the ML model, we used a fixed-length list of numbers representing each *genome_name*, i.e., a feature vector. To featurize the 81 sequenced *genome_names* using the PATRIC annotations, we counted the number of times a particular value occurs in the PATRIC *Superclass, Class, Subclass, Subsystem* name and *Role ID* columns for a given *genome_name*.

For example, if a particular *genome_name* maps to exactly seven PATRIC IDs for which the *PATRIC Subsystem* column’s value is “DNA repair, bacterial,” we create a feature column named “Subsystem Name=DNA repair, bacterial” whose feature value on the row for that *genome_name* is 7.

We further annotated the featurized DataFrame with additional feature columns for each *genome_name* with binary indicator variables for its location within the phylogenetic tree. For example, there was a column named “Family=Bacteroidaceae” whose value was 1 for every *genome_name* in the Bacteroidaceae family and 0 for other genome_names. We added these indicator variables for all observed values in the *Phylum, Class, Order, Family, Genus*, and *Species* columns.

The feature DataFrame is relatively sparse and is thus filled with many cells containing counts of 0, as many *genome_names* did not associate with values for the PATRIC columns whose values we counted, but we did not take advantage of this sparsity.

#### Model output values

We joined the growth data outputs (maximum OD across pH and osmolality conditions, Table S5) for each strain’s genome with the information obtained based on the *genome_name*.

We constructed separate models for predicting pH and osmolality responses. Within those models, we jointly modeled all observations relevant to each perturbation. For example, we constructed a single model that jointly predicts the normalized maximum OD observations across all pH conditions based on the above-described features representing the *genome_name*. Thus, our models for predicting pH response have four real-valued outputs for the observations at pH = 4, 5.4, 6.7, and 7.3. Likewise, the osmolality prediction models have four outputs, for the lowest osmolality, 890 mOsm/kg, 1180 mOsm/kg, and 1800 mOsm/kg.

The osmolality measurements for one of the *genome_names* failed at two osmolality values; hence, we removed this row from the DataFrame used to fit the osmolality models, leaving 79 rows. The pH data had only one failed row.

#### Model architecture, loss function, and training procedure

Our featurized Pandas DataFrame was very short and wide (81 × 11,514), with a single row for each of the 81 different *genome_names* and 11,514 different feature columns. This shape is atypical for ML applications due to the potential for overfitting, but we were interested in feature evaluation rather than precise modeling; thus, we used a one-level decision tree regression model, also known as a decision stump (58). We fit this model using sklearn’s default parameter values for regression trees, which minimizes the total squared error on both sides of the decision stump (58, 59).

#### Decision stump squared-error equation

A decision stump divides a dataset into two groups using a single numerical feature and a splitting threshold. Rows in which the feature value is smaller than the threshold go down the left branch of the tree and arrive at the left terminal leaf node, whereas rows for which the feature value is at least as large as the threshold go to the right terminal leaf node.

At each of the left and right leaf nodes, the fitting algorithm computes the mean value for each of the outputs using the rows assigned to that node, which serves as the model’s prediction for all rows with the same classification. The fitting algorithm attempts to select a feature and splitting threshold which minimizes the total squared error of each row’s distance from its associated mean. Pedregosa et al. describes this algorithm as minimizing the “mean squared error, which is equal to variance reduction as feature selection criterion and minimizes the L2 loss using the mean of each terminal node” (59).

#### Single-feature decision stumps

To identify the quality of each candidate feature, we trained a single-feature decision stump on all 11,514 individual features. Because the decision tree fitting code is not required to select between competing features for the root split, the only remaining task is to identify the splitting threshold that minimizes the total squared error across the left and right terminal leaf nodes in predicting the maximum OD across all four responses. Note that this approach places us in the regime of a decision tree regression multi-output problem (59).

#### K-fold Cross Validation

Even though the single-feature decision stump is a relatively simple model, it is still possible to overfit the data. The large number of features being evaluated increases the chance that this will happen for some features. To mitigate this risk, we performed k-fold cross validation for all the models we fit, using sklearn’s default parameters settings (K=5). K-fold cross validation works by dividing the shuffled data into 5 partitions. Each time a model is fit, one of the partitions is held out of the training set. We then evaluated the trained model’s squared error on both the training partitions and the held-out test set partition.

Because we trained K=5 different models for each feature, we compute 5 different estimates for each feature’s squared error on the both the train and test sets. To compute an overall estimate of each feature’s quality, we took the mean of those 5 estimates.

We note that, particularly in modeling the pH data, some features happen to result in low squared error on the test set, despite having relatively high error on the train set (**Figure S4B**). To further mitigate this, we computed each feature’s rank amongst all the features in the train and test set scores, and ultimately ranked the features according to its maximum (worst) rank across the train and test set scores.

Finally, since we trained 5 different models for each feature, each of the models could have selected a different decision splitting threshold for that feature. For binary partitions of the data used to make box plots and heatmaps, we took the mean value of the decision threshold selected for each of the 5 models.

### Colony polymerase chain reaction and confirmation of strains used in experiments

Here, 1 mL of overnight cultures grown for 24 hours at 37°C was spun down for 5 minutes at 5000 x g (relative centrifugal force), and the collected pellets were boiled for 5 minutes. We prepared 1:10 and 1:100 dilutions using sterile DNase-free water to prepare for colony polymerase chain reaction (PCR) ([98°C 2 min] → [98°C 30 s, 57°C 30 s, 72°C 45 s] x 30 cycles → [72°C 10 min] → [4°C ∞]) and performed sequencing using 8F and 1391R primers (8F: AGAGTTTGATCCTGGCTCAG, 1391R: GACGGGCGGTGWGTRCA). Some cultures required DNA extraction prior to PCR, which was performed using the Qiagen DNeasy Blood and Tissue Kit (Catalog Number: 69504).

### Human fecal sample collection, fermentation, and DNA extraction

We collected feces from six individuals (TL1, TL2, TL3, TL4, TL5, TL6), either a day prior to or on the morning of experimentation. We stored all fecal samples in sterile conical tubes at - 80°C before processing. Fecal samples (1.5 g) were resuspended in sterile pre-reduced 1X PBS, and contents were allowed to settle to collect liquid. Liquid Mega Medium was pre-reduced in an anaerobic chamber environment with 5% CO_2_, 5% H_2_, and 90% N_2_ for at least 24 hours. For each sample, 1 μL of supernatant was collected and subsequently inoculated into 200 μL of prereduced liquid Mega Medium adjusted to a pH of 4, 5.5, 6.9, or 7.6 and an osmolality of 472, 670, 862, 1047, 1247, 1437, 1637, or 1824 mOsm/kg on two 96-well plates. We measured the OD at a wavelength of 600 nm (OD_600_) using a BioTek Synergy H1 Plate Reader after anaerobic incubation at 37°C for 48 hours. We selected the physical conditions based on generated OD_600_ measurements and extracted DNA from the 96-well plates using the DNeasy PowerSoil Pro Kit (Catalog Number: 47016).

### Humanized mice supplemented with a guar gum diet

Germ-free Swiss Webster (SW) mice were gavaged with a human gut microbiota (TL1) at 9 weeks and placed on a standard rodent diet (LabDiet 5k67). Six weeks after colonization, 5 SW mice (2 males, 3 females) were switched to a guar gum diet (TestDiet 5BSE) for 2 weeks while 3 (3 males) remained on the standard diet. Two weeks after equilibration on the diet, the mice were sacrificed using carbon dioxide with secondary cervical dislocation. We collected contents from the duodenum, jejunum, ileum, cecum, and colon in the gastrointestinal tract for pH and osmolality measurements. Collected mouse intestinal contents were stored in 1.5-mL microcentrifuge tubes and kept on ice during preparation for pH and osmolality measurements. The same micro pH probe (Orion™ PerpHecT™ ROSS™ Combination pH Micro Electrode, Cat. No. 8220BNWP) and Advanced Instruments Osmo1® Single-Sample Micro-Osmometer were used to measure intestinal contents as described above. In addition, we performed DNA extraction as described above for 16S rRNA sequencing and sent cecal contents for SCFA analysis.

### 16S sequencing library preparation and sequencing

We quantified the extracted DNA using Quant-iT 1X dsDNA HS (High-Sensitivity) and Quant-iT 1X dsDNA BR (Broad Range) Assay kits (Catalog Number: Q33232). We submitted the DNA samples to either Biofactorial, a high-throughput biology facility located in the Life Sciences Institute at the University of British Columbia, or the Gut4Health Microbiome Core Facility at the British Columbia Children’s Hospital Research Institute and the University of British Columbia for a paired-end sequencing run using a MiSeqv3-600 instrument with dual-indexed V4V5 primers (Biofactorial) or V4 primers (Gut4Health). For samples sequenced at Biofactorial, a dual-indexing, one-step 10-μL PCR reaction was performed on a LabCyte Access Workstation using Quanta repliQa HiFi ToughMix with 0.5 ng input DNA and complete “fusion primers” that include Illumina Nextera adaptors, indices, and specific regions targeting the V4/V5 region of the16S rRNA genes (60). After quantification of amplicons via a picogreen assay (Quant-iT™ PicoGreen™ dsDNA Assay Kit, Thermo Fisher), 2 ng of each product were pooled for subsequent cleanup using the AmpureXP PCR cleanup protocol (Beckman). The pooled library was quantified by a picogreen assay and loaded onto an Illumina MiSeq Reagent Kit v3 (600-cycle) according to the manufacturer’s recommendations with 15% PhiX. Samples submitted to Gut4Health were prepared according to a previously published method (61). Briefly, the V4 region of the 16S rRNA gene was amplified with barcode primers containing the index sequences using a KAPA HiFi HotStart Real-time PCR Master Mix (Roche). We monitored PCR product amplification and concentration on a Bio-Rad CFT Connect Real-Time PCR system. Amplicon libraries were then purified, normalized, and pooled via a SequalPrep™ normalization plate (Applied Biosystems). We further purified the pooled library with the Agencourt AMPure XP system (Beckman Coulter) according to the manufacturer’s protocol. Library concentrations were verified by a Qubit™ dsDNA high-sensitivity assay kit (Invitrogen) and the KAPA Library Quantification Kit (Roche) according to the manufacturers’ instructions. We submitted the purified pooled libraries to the Bioinformatics + Sequencing Consortium at the University of British Columbia, which verifies DNA quality and quantity using a high-sensitivity DNA kit (Agilent) on an Agilent 2100 Bioanalyzer. Sequencing was performed on the Illumina MiSeq™ v2 platform with 2 × 250 paired end-read chemistry.

### 16S rRNA data analysis

We imported 16S rRNA reads generated through the MiSeq v3-600 run to QIIME 2 2020.11 for analysis (62). The read quality was assessed by FastQC (https://github.com/s-andrews/FastQC). We processed the sequences using DADA2 (https://github.com/qiime2/q2-dada2). The primers were trimmed, and sequences with a quality score below 30 were truncated. We taxonomically classified the denoised sequences using a SILVA v138-trained classifier (63).

### SCFA analysis of murine cecal samples from the guar gum diet

To prepare SCFA samples for gas chromatography analysis, we extracted SCFAs from 40–100 mg of cecal contents. The samples were mixed and homogenized with 800 μL of 25% phosphoric acid. We collected the supernatants via centrifugation at 15,000 x g for 10 minutes at 4°C. We then added 200 μL isocaproic acid and 0.2 mL 25% phosphoric acid as an internal standard. Supernatants were sent to the AFNS Chromatography Facility at the University of Alberta for quantification. We ran the samples on a Varian 430 gas chromatograph with a Stabilwax-DA column (length: 30 m, inner diameter: 0.53 mm, film thickness: 0.5 μm) and helium carrier gas, using a 250C injector with a split ratio of 5 and a 1-μL injection. We utilized a flame ionization detector with a detector temperature of 250°C. Retention times were compared with known standards.

### Software and algorithms

In this work, we utilized MATLAB 2020a (MathWorks) to analyze the bacterial growth, as described above. We used BioTek GEN5 software to collect bacterial absorbance and fluorescence. We employed RStudio to compare the abundance of annotated genomic features across species at subsystem levels (Shiny library) in RStudio. We performed analyses and graphing using R V4.1.2 and RStudio V1.4.1717, as described above.

## Acknowledgments

The authors acknowledge that the land we performed this research on is the traditional, ancestral, and unceded territory of the xwməθkwəýəm (Musqueam) Nation. We encourage others to learn more about the native lands in which they live and work at https://native-land.ca/. The authors acknowledge support from the Canadian Institutes of Health Research Team Grant: Canadian Microbiome Initiative 2 (to all), Crohn’s and Colitis Canada (to all), Canadian Institute for Advanced Research (to all), Michael Smith Foundation for Health Research Scholar Award (to C.T.), Paul Allen Distinguished Investigator Award (to C.T.), the University of British Columbia Killam Postdoctoral Fellowship (to J.N.), a National Science Foundation Postdoctoral Fellowship in Biology (to J.N.), and the Johnson & Johnson WiSTEM2D Award (to C.T.). This work also received support from the Life Sciences Institute Biofactorial High-Throughput Biology Core, supported by the University of British Columbia Global Research Excellence Biological Resilience Initiative. We also acknowledge Dr. KC Huang for useful discussions, Dr. Lisa Osborne and Naomi Fettig for support in providing special diets for our mouse experiments, Negin Rahanjam for support in animal husbandry, and Dr. Ho Pan Sham and Dr. Catherine Chan (Gut4Health) and Dr. Tom Pfeifer (Biofactorial) for assistance with next-generation sequencing. We thank Dr. John Nomellini for research support and Dr. Richa Anand and Dr. Nina Maeshima for providing editorial support for grants that ultimately funded this work. We also thank Giselle McCallum, Sophie Cotton and Dr. Lisa Osborne for critically reading this manuscript and providing feedback.

## Supplementary Material Legends

**Figure S1. pH changes due to fermentation do not correlate with growth.** A) Maximum OD versus change in pH (measured with BCECF; Materials and Methods) under different conditions, labeled by bacterial family. B) Maximum OD versus maximum growth rate under different conditions, labeled by bacterial family.

**Figure S2. Osmolality (left) and pH (right) features show varying degrees of correlation.** Each square represents the number of strains for which the two thresholded features have the same value.

**Figure S3. Osmotic features identified by ML analysis are generalizable.** A) Measure of ML model generalization by comparing performance of model on training and held out test sets. Each point is a PATRIC feature for which we evaluated 5 K-fold models. B-C) Phenotypic variations in response to osmotic stress correlated with cold shock proteins: correlations occur both *in vitro* (B) and *in vivo* (C).

**Figure S4. Normalized maximum OD at pH 5.4 versus total squared error for each feature.** A) Most PATRIC features identified with ML predict that the presence of a specific feature will lead to a lower maximum normalized OD at pH 5.4. For features with a normalized maximum OD closer to 1, indicating good growth in acidic conditions, the features are heavily correlated with the family Lactobacillaceae (Materials and Methods). B) Measure of ML model generalization by comparing performance of model on training and held out test sets. Each point is a PATRIC feature for which we evaluated 5 K-fold models.

**Figure S5. PATRIC features do not correlate in their ability to predict growth in different osmolality and pH conditions.** Total squared error for osmolality conditions versus total squared error for pH conditions.

**Table S1. Medium components for growth media used in the study.** Details are given for each strain, including the medium components, source, and sequencing results.

**Table S2. PATRIC features identified by ML to distinguish growth in different osmolality conditions.** Columns indicate the name of the PATRIC feature, the threshold on the PATRIC count of the feature that distinguishes the phenotype, the number of strains whose count is lower than the threshold number of the feature, the number of strains whose count is higher than the threshold, and the total squared error for the mean of all ODs (Materials and Methods).

**Table S3. PATRIC features identified by ML to distinguish growth in different pH conditions.** Columns indicate the name of the PATRIC feature, the threshold on the PATRIC count of the feature that distinguishes the phenotype, the number of strains whose count is lower than the threshold number of the feature, the number of strains whose count is higher than the threshold, and the total squared error for the mean of all ODs (Materials and Methods).

**Table S4. PATRIC features used in the ML model.** Each strain in our collection was annotated with PATRIC, and all features are represented in this table.

**Table S5. Normalized growth rate (GR) and maximum OD for all sequenced strains used in the ML model under different growth conditions.** Growth data outputs for each strain’s genome are combined with information obtained based on the *genome_name*.

## Notes

### Competing Interest Statement

The authors have declared no competing interest.

