## Supplementary figures and images for "Single-strain behavior predicts responses to environmental pH and osmolality in the gut microbiota"

### Supplemental Figure 1

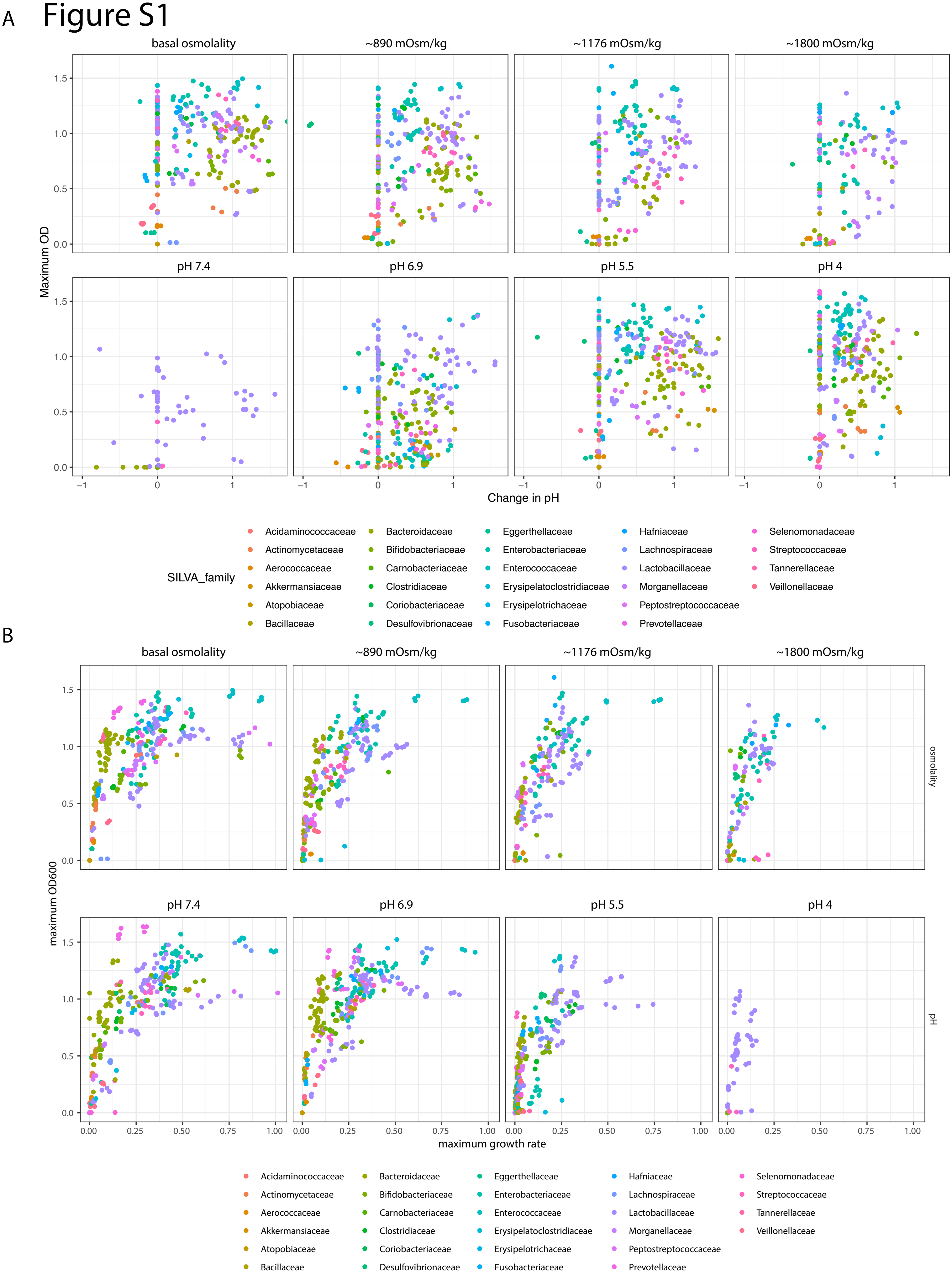

### Supplemental Figure 2

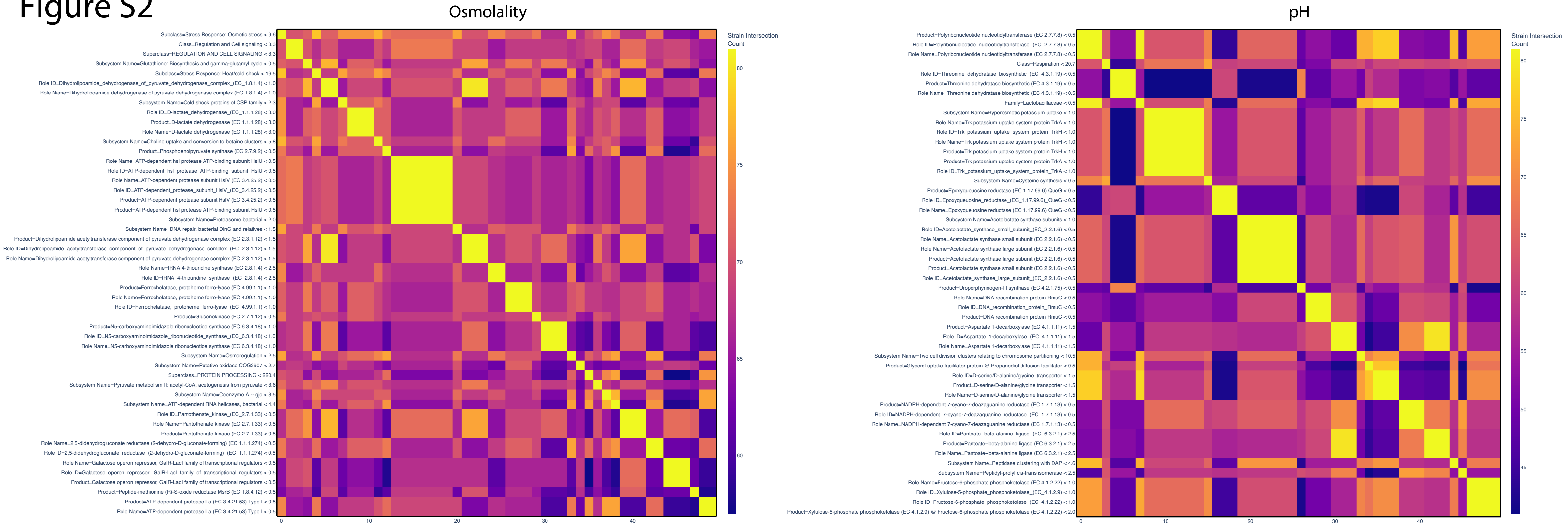

### Supplemental Figure 3

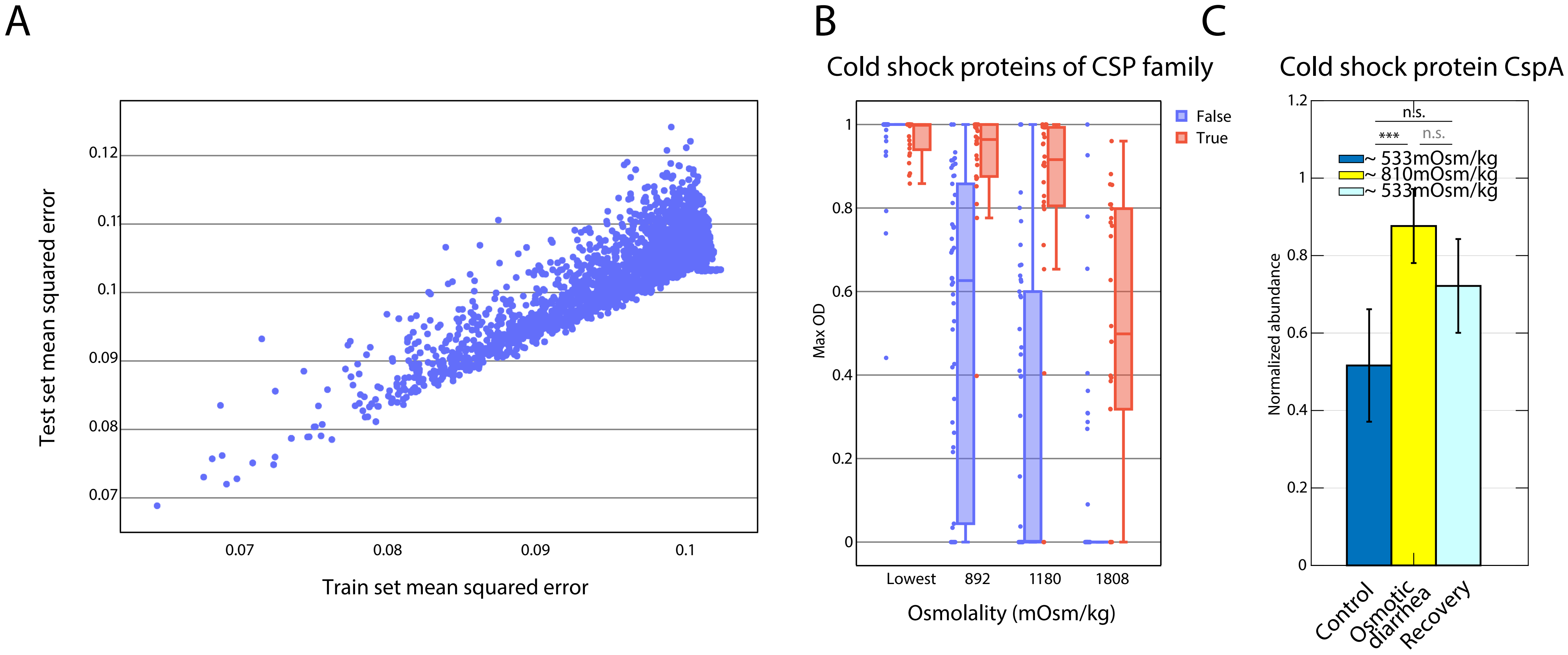

### Supplemental Figure 4

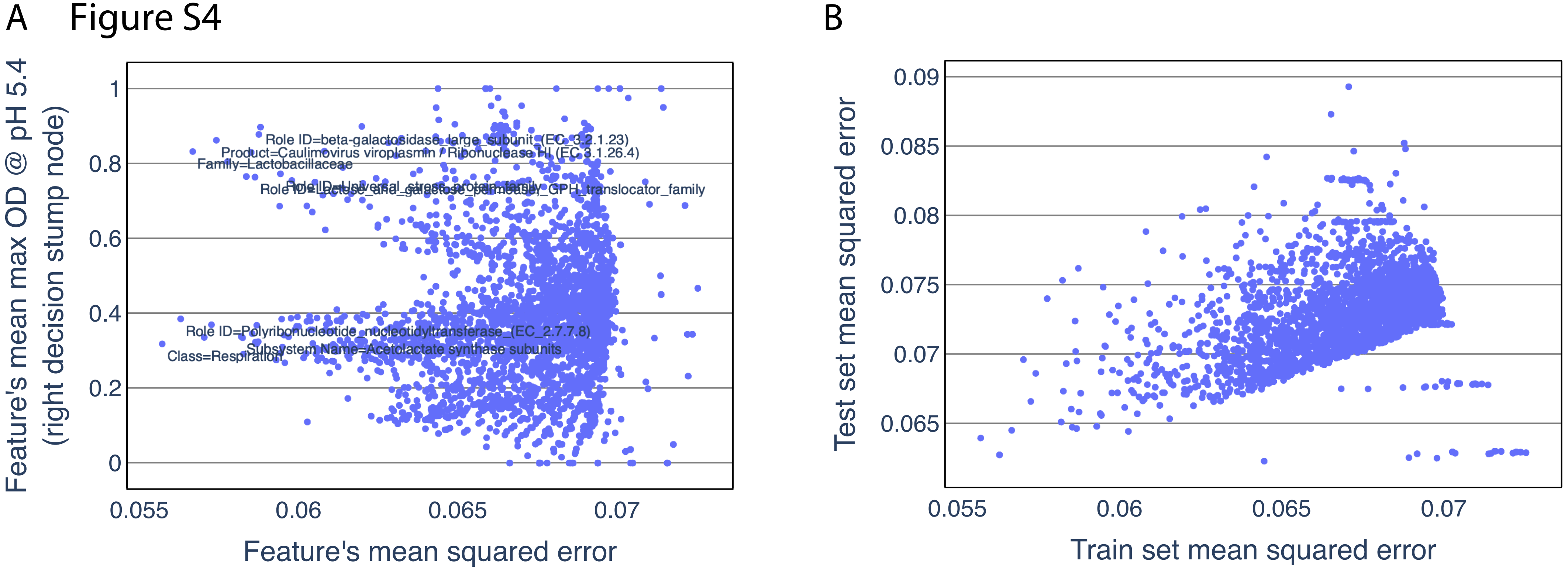

### Supplemental Figure 5

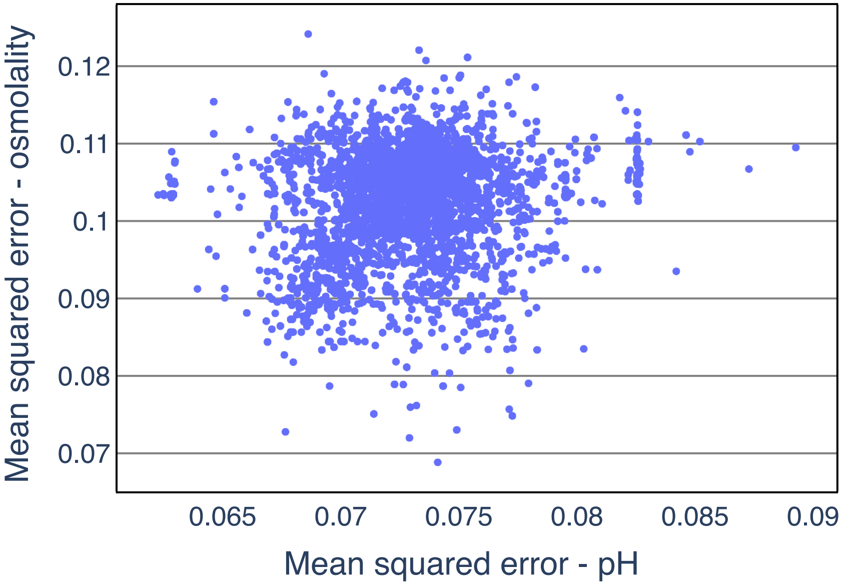
